# Transport of herbicides by PIN-FORMED auxin transporters

**DOI:** 10.1101/2024.08.29.610046

**Authors:** Lukas Schulz, Kien Lam Ung, Sarah Koutnik-Abele, David L Stokes, Bjørn Panyella Pedersen, Ulrich Z. Hammes

**Author notes:** These authors contributed equally to this work (Lukas Schulz, Kien Lam Ung). To whom correspondence should be addressed. (B.P.P.); (U.Z.H.).

## Abstract

Auxins are a group of phytohormones that control plant growth and development ^1^. Their crucial role in plant physiology has inspired development of potent synthetic auxins that can be used as herbicides ^2^. Phenoxyacetic acid derivatives are a widely used group of auxin herbicides in agriculture and research. Despite their prevalence, the identity of the transporters required for distribution of these herbicides in plants is both poorly understood and the subject of controversial debate ^3,4^. Here we show that PIN-FORMED auxin transporters transport a range of phenoxyacetic acid herbicides across the membrane and we characterize the molecular determinants of this process using a variety of different substrates as well as protein mutagenesis to control substrate specificity. Finally, we present Cryo-EM structures of *Arabidopsis thaliana* PIN8 with 2,4-dichlorophenoxyacetic acid (2,4-D) or 4-chlorophenoxyacetic acid (4-CPA) bound. These structures represent five key states from the transport cycle, allowing us to describe conformational changes associated with substrate binding and transport across the membrane. Overall, our results reveal that phenoxyacetic acid herbicides use the same export machinery as endogenous auxins and exemplify how transporter binding sites undergo transformations that dictate substrate specificity. These results enable development of novel synthetic auxins and for guiding precision breeding of herbicide resistant crop plants.

## INTRODUCTION

Auxins are a group of plant hormones that regulate development and growth responses in plants. These hormones are distributed through a process known as Polar Auxin Transport (PAT), which underlies fundamental environmental responses such as gravitropic and phototropic growth, root development and lateral root initiation, as well as organ initiation and differentiation ^5–7^. PIN-FORMED (PIN) auxin transporters are key for providing polarity to auxin transport by mediating auxin export from the cytosol to the apoplast ^8,9^. The principal endogenous auxin is indole-3-acetic acid (IAA, *pK_a_* = 4.7), and a variety of synthetic auxins are used as herbicides, together accounting for ∼20% of all herbicide-treated farmland world-wide ^10^. In general, the synthetic auxins elicit typical auxin responses, but with stronger effects that are due in part to longer half-lives in the cytosol ^2,10–12^. A major subgroup of synthetic auxins comprises phenoxyacetic acids that include the widely used 2,4--dichlorophenoxyacetic acid (2,4-D, *pK_a_* = 2.73) and 4-chlorophenoxyacetic acid (4-CPA, *pK_a_* = 3.56). Despite their high agricultural and ecological impact, it remains unclear how these compounds are distributed across plant tissues. In particular, it remains controversial if PINs are the responsible exporters and this ambiguity makes it impossible to understand how the observed chemical diversity of synthetic auxins is tolerated ^2,13^.

Recently, structures of *Arabidopsis thaliana* PIN8, PIN1 and PIN3 have been determined, revealing symmetric homodimers, each of which harbour ten transmembrane helices (M1-M10) with an inverted repeat that produces two 5-helix bundles ^14–17^. The first two helices from each repeat (M1-M2 and M6-M7) form a scaffold domain that mediates dimerization, and a long cytoplasmic loop between M5 and M6 that connects the two repeats. A transport domain is composed of M3-M5 and M8-M10, also related by the inverted symmetry, and a substrate binding chamber is centred on the crossover of M4 and M9, which form interrupted α-helices ^15,16^. PIN8 produced structures in both inward and outward conformations, thus showing that PINs operate by an elevator mechanism where the substrate-loaded transporter domain is translated 5 Å across the membrane relative to the scaffold domain ^16,18^. In the inward open conformation, the binding chamber connects to a larger vestibule leading to the cytosol, whereas in the outward open conformation the binding chamber connects directly to the extracellular space ^15,16^. All existing structures show symmetric dimers in which monomers adopt identical conformations ^14–17^.

Here, we address phenoxyacetic acid herbicide transport and broader implications for substrate recognition in PINs. We show that PINs can bind and transport these synthetic auxins *in vitro* and that these substrates are distributed *in planta* in a PIN-dependent fashion. In addition, we characterize binding and transport properties of a variety of related natural and synthetic auxins *in vitro* and we use single particle cryoEM to solve structures of PIN8 in complex with 2,4-D or 4-CPA to confirm their role as PIN substrates and to clarify how the chemical diversity of synthetic auxins can be tolerated in the binding site. In the case of 4-CPA, a single sample produced five different structures representing key binding intermediates from the transport cycle and indicate that PIN monomers operate asynchronously. Together with the biochemical characterization the structures pinpoint key elements of PIN proteins and auxin substrates that control affinity and specificity. This information may be useful in engineering herbicide resistant crops as well as novel environmentally safer herbicides for sustainable agriculture.

## RESULTS

Trans-cellular transport and distribution of 2,4-D *in planta* has been shown to be active, polar and sensitive to IAA export inhibitors, but much slower than IAA transport ^4,19,20^. These results suggested that 2,4-D is transported by PIN proteins. In contrast, one influential study suggested that the low rate of 2,4-D export implies passive diffusion across the membrane, rather than a protein-mediated transport process ^3^. To investigate this discrepancy and understand the basis for auxin herbicide transport, we used solid supported membrane (SSM) electrophysiology to characterize 2,4-D binding and transport by purified *Arabidopsis thaliana* PIN8 reconstituted into proteoliposomes (Fig. 1A). We find that the peak current responses elicited by 2,4-D and IAA are similar (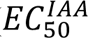 250 ± 10 µM; 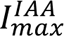 18 ± 3 nA vs 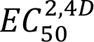 180 ± 10 µM; 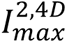17 ± 2 nA) (Fig. 1A). The current response to 2,4-D, like that of IAA, is inhibited by the PIN-inhibitor naphtylphtalamic acid (NPA), suggesting competitive binding (Fig. 1A inset). Based on this inhibition, we determine the 2,4-D binding affinity (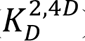) to be 60 ± 10 µM which is almost identical to the previously described IAA binding affinity (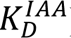) of 40 ± 15 µM (Extended Data Fig. 1A and D, and ^16^). These *in vitro* PIN8 results are corroborated by *Xenopus* oocyte efflux assays using both PIN8 and PIN3 (Fig. 1B and C). These assays show that PIN8 mediates NPA--sensitive 2,4-D export from oocytes with a transport rate of about half of that determined for IAA (Fig. 1B and Extended Data Fig. 2A-C). To show that 2,4-D export is not unique to PIN8, we tested *A. thaliana* PIN3 with and without the activating PINOID (PID) kinase ^21^. As expected, PIN3 is activated by kinase and, as with PIN8, the NPA-sensitive, 2,4-D export rate by PIN3 is about half of the IAA transport rate (Fig. 1C and Extended Data Fig. 2D-F). To test directional transport in plants we used an *A. thaliana* stem segment assay and compared 2,4-D and IAA transport in stem segments of wild type and *pin1* mutants since PIN1 is the transporter responsible for auxin transport in this organ (Extended Data Fig. 2G-I) ^21^. This assay shows that although 2,4-D transport is about 20-fold lower than IAA transport in wildtype plants, it is completely abolished in *pin1* mutants, demonstrating that 2,4-D transport *in planta* is in fact PIN-mediated.

**Fig. 1:**
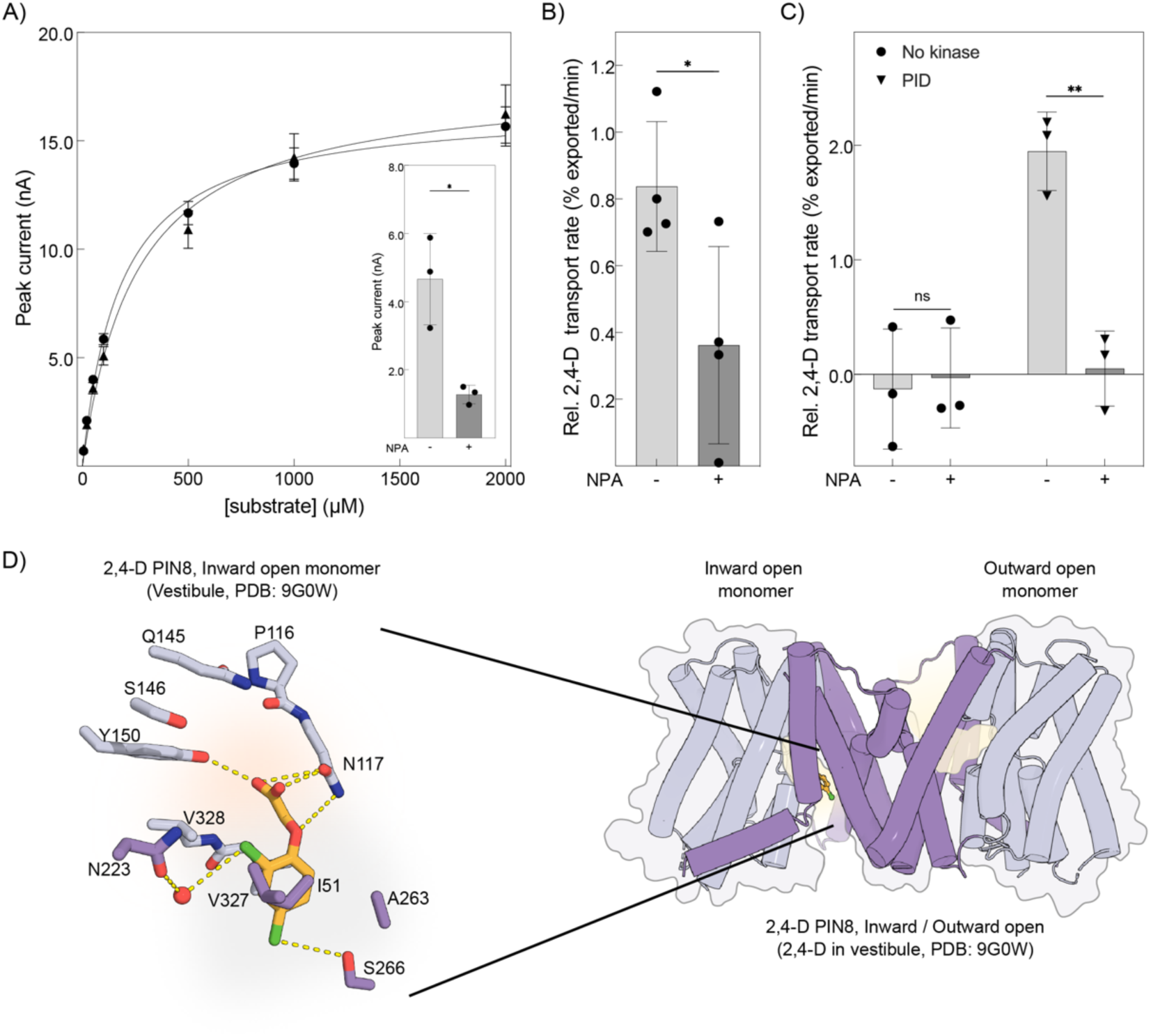
2,4D is a PIN substrate. A) Peak current response as a function of [IAA] (▴) and [2,4D] (●) determined by SSM electrophysiology on PIN8 proteoliposomes. A Michaelis–Menten model is fit to describe kinetics (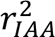 = 0.9579; 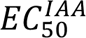 254 ± 6 μM; 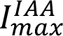18 ± 3 nA vs 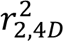 = 0.9802; 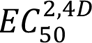 179 ± 10 μM; 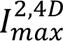 17 ± 2 nA; data points represent mean ± s.e.m.; n = 4 individual sensors). Inset: Peak current response elicited by 100μM 2,4D in absence or presence of 20 µM NPA. Data are mean ± s.e.m. of n = 3 individual sensors. Groups were compared by a two-sided unpaired Student’s t-test. p = 0.0126. B) Relative transport rates of [3H]-2,4D export by PIN8 in presence or absence of 10 μM NPA as indicted from oocytes calculated from four independent time course experiments. Data points are individual experiments and bars represent mean ± SD. Groups were compared by a two-sided unpaired Student’s t-test P = 0.0361; n = 4 individual experiments. C) Relative transport rates of [3H]-2,4D export by PIN3 with or without the activating PID kinase in presence or absence of 10 μM NPA as indicted from oocytes calculated from three independent time course experiments. Data points are individual experiments and bars represent mean ± SD. Groups were compared by a two-sided unpaired Student’s t-test. n.s. not significant P = 0.6411; ** P = 0.0004; n = 3 individual experiments. D) Structure of asymmetric dimeric PIN8 with 2,4-D bound to the inward open monomer in a prebinding position in the vestibule (right panel). The transporter domain highlighted in light violet and the scaffold domain in violet and ligand coloured in orange. Close-up view of 2,4-D and the residues from the scaffold domain (violet) and transporter domain (light violet) interacting with the substrate. The inward-open pocket is divided into the binding chamber (highlight in light orange) and a vestibule (highlight in grey). Red spheres represent water molecules.

To supplement the transport data, we use cryo-EM to determine the structure of PIN8 bound to 2,4-D at 3.5 Å resolution (Fig. 1D, Extended Data Fig. 3 and 4A, Extended Data Table 1). Notably, the homo-dimer structure is asymmetric where one monomer adopts an inward facing (IF) conformation and the other an outward facing (OF) conformation. The monomer in the inward conformation shows 2,4D bound within the vestibule seen in previous structures, with a pose that suggests a prebinding position. In particular, the carboxyl-group of 2,4D interacts with Asn117 and interrupted helices associated with the crossover (M4 and M9) are positioned such that their dipoles neutralize the negative charge. In contrast, the monomer in the outward facing conformation has weak density in the binding pocket that cannot be confidently modelled, but that appears consistent with 2,4-D release (Fig. 1D).

To explore chemical determinants of phenoxyacetic acid herbicides and to define crucial interactions within the binding chamber of PIN8, we measured binding affinity (*K_D_*) for a set of derivative compounds based on their competition with the inhibitor NPA. Specifically, we used the SSM assay to assess inhibition of the strong binding current produced by NPA [Extended Data Fig. 1 and Extended Data Fig. 4 A and B and ^16^]. To start, we analysed the known herbicides 2,4,5-trichlorphenoxyacetic acid (2,4,5-T), 2-methyl-4-chlorphenoxyacetic acid (MCPA) and 4-chlorophenoxyacetic acid (4-CPA) in addition to 2,4-D, all of which share three critical features: the carboxylate group, the aryl ether group, and a 4-Cl group (Fig. 2A and B, dark grey bars). Compared to IAA we find that 4-CPA binds with significantly lower affinity (∼8-fold, p < 0.05) (Fig. 2A and Fig. S1A and S1I) confirming recent results for this compound (12). 2,4 D binds with similar affinity to IAA, whereas MCPA binds with intermediate affinity (Fig. 1A, Fig. 2A and Extended Data Fig. 1A, 1D, 1E, 1F 1I). 2,4,5-T binds with the highest affinity of all the substrates tested (Fig. 2A and Fig. S1B). These results show that the number of substitutions in the phenyl ring, their position, chemical identity and influence on the π-orbital electrons impact substrate binding.

**Fig. 2:**
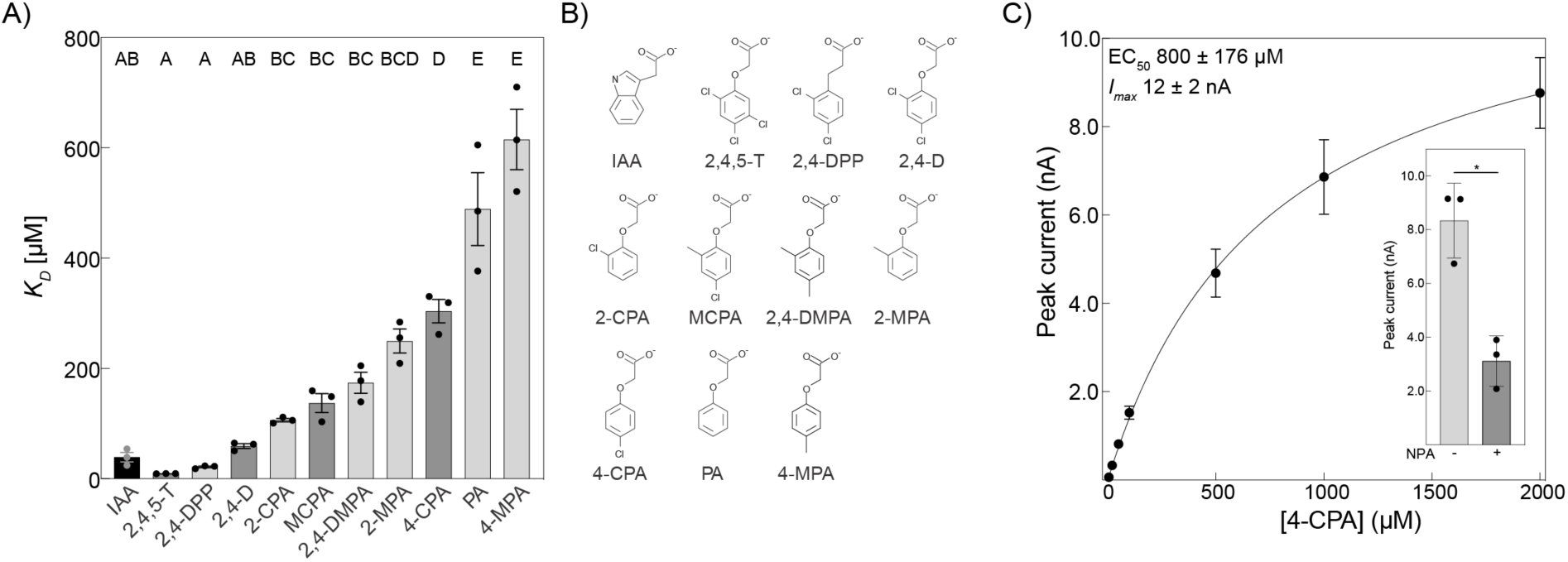
Substrate determinants and 4-CPA binding affinity. A) *K_D_* of substrate binding by PIN8 as determined by SSM electrophysiology. Dark grey represents substrates with herbicidal activity. Indole-3-acetic acid (IAA); 4-Chlorphenoxyacetic acid (4-CPA); 2,4-Dichlorphenoxyacetic acid (2,4D); 2,4,5-Trichlorphenoxyacetic acid (2,4,5-T); 4-Chloro-2-methylphenoxyacetic acid (MCPA); Phenoxyacetic acid (PA); 2-Chlorphenoxyacetic acid (2-CPA); 2-Methylphenoxyacetic acid (2-MPA); 3(2,4-dichlorphenyl)-propionic acid (2,4-DPP). Data points are individual sensors and bars represent mean ± s.e.m. Data points derived from the full measurements (Extended Data Fig. 1). Letters indicate significance differences between groups determined by a one-way ANOVA followed by a Turkeýs multiple comparisons test (p < 0.05). For individual p-values see Extended Data Table 3. B) Chemical structures of the substrates tested are shown at pH 7.4. C) Peak current response as a function of [4-CPA] determined by SSM electrophysiology on PIN8 proteoliposomes. A Michaelis–Menten model is fit to describe kinetics (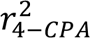 = 0.98; 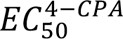 800 ± 200 μM; 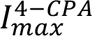 12 ± 2 nA; data points represent mean ± s.e.m.; n = 3 individual sensors). Inset: Peak current response elicited by 2 mM 4-CPA in absence or presence of 20 µM NPA. Data are mean ± s.e.m. of n = 3 individual sensors. Groups were compared by a two-sided unpaired Student’s t-test. P = 0.0057.

To characterize the impact of substitutions in the phenyl ring on substrate affinity we investigated several compounds displaying substitutions at different positions and with different chemistries. We find that phenoxyacetic acid (PA), which lacks any substitutions at the phenyl ring, binds with significantly lower affinity (*K_D_* ∼500 µM) (Fig 2A and B and Extended Data Fig. 1J), indicating that chloride substitutions in the ring increase substrate affinity. Next, we determine that 3-(2,4-dichlorphenyl)-propionic acid (2,4-DPP) binds with higher affinity than 2,4-D suggesting that the aryl ether is not important (Fig 2A and B and Extended Data Fig. 1C and D). 2-chlorophenoxyacetic acid (2-CPA), is bound with high affinity, comparable to that of IAA and 2,4-D (Fig 2A and B and Extended Data Fig. 1A, D and E), suggesting that substitution at the C2 position on the phenyl-ring is more important than at the C4 position (c.f., 2-CPA vs. 4-CPA). Specificity for *chloride* substitutions is evaluated by testing 2--methylphenoxyactetic acid (2-MPA, with a methyl group at C2) 4--methylphenoxyactetic acid (4-MPA, with a methyl group at C4) and 2,4--methylphenoxyactetic acid (2,4-DMPA, with a methyl group at C2 and C4) (Fig. 2A and B and Extended Data Fig. 1G, H and K). Corresponding *K_D_* values indicate that methyl groups produce lower affinity than chloride. Methyl groups in both C2 and C4 positions produce higher affinity than at C2 alone, whereas a methyl group at C4 alone produces even lower affinity than PA. This again shows that the C2 position is more important for substrate affinity than C4.

The ability of PIN8 to adopt different conformations during structure determination with 2,4-D suggested it would be possible to capture the key conformations representing a full transport cycle in a single cryo-EM sample. For this, we tried 4-CPA as a substrate which was recently found to be an important member of this group of herbicides with very low affinity for PIN-mediated export ^12^. SSM measurements indicate that 4-CPA elicits NPA-sensitive current responses in PIN8 that have higher *EC_50_* (800 ± 200 µM) and *K_D_* (200 ± 100 µM), consistent with a lower affinity for 4-CPA relative to 2,4-D and IAA (Fig. 2C, Extended Data Fig. 1A, D and I). To establish that 4-CPA is also transported by PIN8 we measured the width of current SSM-based responses at various lipid-to-protein-ratios (LPR) of IAA, 2,4-D and 4-CPA to that of the non-transported inhibitor NPA (Extended Data Fig. 5A and C) ^22^. For IAA, 2,4D and 4-CPA, the peak widths vary with LPR, consistent with transport of these compounds, whereas peak width does not change for NPA, consistent with an inhibitor that produces a binding current but is not transported ^22^.

Analysis of cryo-EM images of PIN8 with 4-CPA produced three independent structures that reveal five distinct conformations coexisting within a single sample (Extended Data Fig. 6, Extended Data Fig. 7A and Extended Data Table 1). These structures comprise both symmetric (OF/OF) and asymmetric (OF/IF) dimers with individual monomers adopting a range of different conformations representing both 4-CPA bound and empty states. By focusing on the ligand binding site for 3D classification, we are able to resolve five unique states: inward-facing empty (IF_empty_ at 3.4 Å), inward-facing with 4-CPA in a prebinding site in the vestibule (IF_4CPA-prebinding_ at 3.4 Å), outward-facing with 4-CPA in the binding pocket (OF_4CPA-bound_ at 3.3 Å), outward-facing with 4-CPA in a partly released state (OF_4CPA-partlyrelease_ at 3.3 Å) and outward-facing empty (OF_empty_ at 3.3 Å) (Fig. 3A-C, Extended Data Fig. 4B-D and Supplementary Movie 1).

**Fig. 3:**
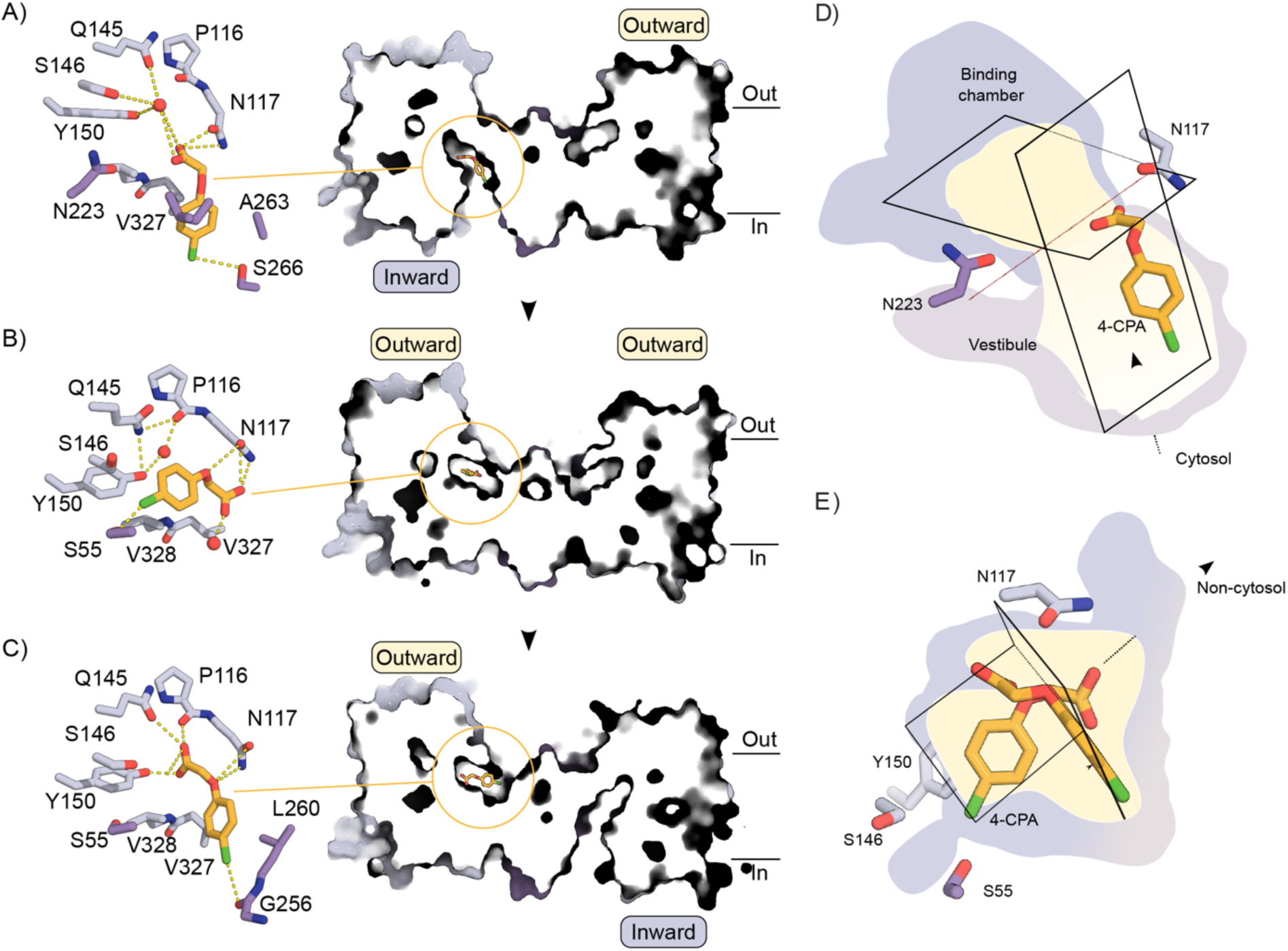
4-CPA structure and transport cycle. A) Close-up view (left) and cut away view of surface presentation (right) of the asymmetric PIN8 structure bound 4-CPA in the prebinding state of the inward monomer. B) Close-up view (left) and cut away view of surface presentation (right) of the symmetric PIN8 structure bound 4-CPA in the binding state of the outward monomer. C) Close-up view (left) and cut away view of surface presentation (right) of the asymmetric PIN8 structure bound 4-CPA in the partly release state of the outward monomer. Residues belonging to the transporter domain highlighted in light violet and the scaffold domain in violet. Ligands coloured in orange. D) Overlay of 4-CPA prebinding position (PDB 9G0X and EMD-50951) in the inward monomer. The inward-facing pocket is divided into the binding chamber (light violet) and a vestibule (grey). The red broken line running from Asn117 to Asn223 defines the boundary between vestibule to binding chamber. E) Overlay of 4-CPA binding (PDB 9G0Z and EMD-50952) and partly release (PDB 9G10 and EMD-50953) position in the outward monomer. Both structures are superposed on the scaffold domain. The superposition shows that in either state, Asn117 retains its interaction with carboxyl and arylether group.

These structural data reflect the transitions of PIN8 and its substrate during the transport cycle (Fig. 3A-C and Extended Data Fig. 4B-D). In IF_4CPA-prebinding_, 4-CPA is located in the vestibule of PIN8 in a position similar to 2,4-D as described above: the carboxyl group is coordinated by Asn117, while a water molecule mediates contact to Gln145 and Tyr150 in the binding chamber. The 4-Cl atom is coordinated by Ser266 (Fig. 3A and Extended Data Fig. 4B). OF_4CPA-bound_ shows 4-CPA fully engaged with the binding chamber (Fig. 3B and Extended Data Fig. 4C). Here, the carboxyl group of 4-CPA is coordinated by Asn117 and its charge is stabilized by the two dipoles of the crossover, the 4-Cl atom is now coordinated by Ser55. It is notable that the phenyl ring in OF_4CPA-bound_ is bound deeper in the binding chamber relative to IF_4CPA-prebinding_, suggesting that substrate recognition is initiated by binding of the carboxylate group followed by flipping and nestling of the phenyl ring into this chamber (Fig. 3D). OF_4CPA-partlyrelease_ shows that steps of substrate release are similar to those of binding, but in reverse. In the transition from OF_4CPA-bound_ to OF_4CPA-prerelease,_ the phenyl ring is solvated and flips out of the chamber while the carboxyl maintains its interaction with Asn117 (Fig. 3C and E and Extended Data Fig. 4D). Together, these five structures allow us to follow the substrate as it transits through PIN8, highlighting various substrate poses and their transient and changing key interactions with the protein during transport.

To identify molecular determinants of PIN8 substrate recognition, we examined the IF_4CPA-prebinding_ structure to identify key residues and then used SSM electrophysiology to compare binding of 4-CPA, IAA and 2,4-D to single-site mutants (Fig. 4 and Extended Data Fig. 4B and C). It is well-established that Asn117 is critical for coordination of the carboxylate during IAA transport ^16^. Accordingly, the N117A mutation abolishes the current response for both 2,4-D and 4-CPA (Extended Data Fig. 5D). Ser146 and Tyr150 interact with the carboxylate via a water molecule (Fig. 1D and 3A and ^16^). Consistent with this indirect coordination, S146A and Y150F mutations reduce substrate affinity slightly (1.8- to 4.6-fold) while Y150A reduce substrate affinity further (3.3- to 4.8-fold) for all three compounds. Introduction of a bulky side chain with the I51F mutation reduces substrate affinity, likely by hindering substrate entry into the prebinding state. Ser266 directly coordinates the 4-Cl residue (Fig. 1D and 3A) and the S266A mutation reduces affinity for all substrates.

**Fig. 4:**
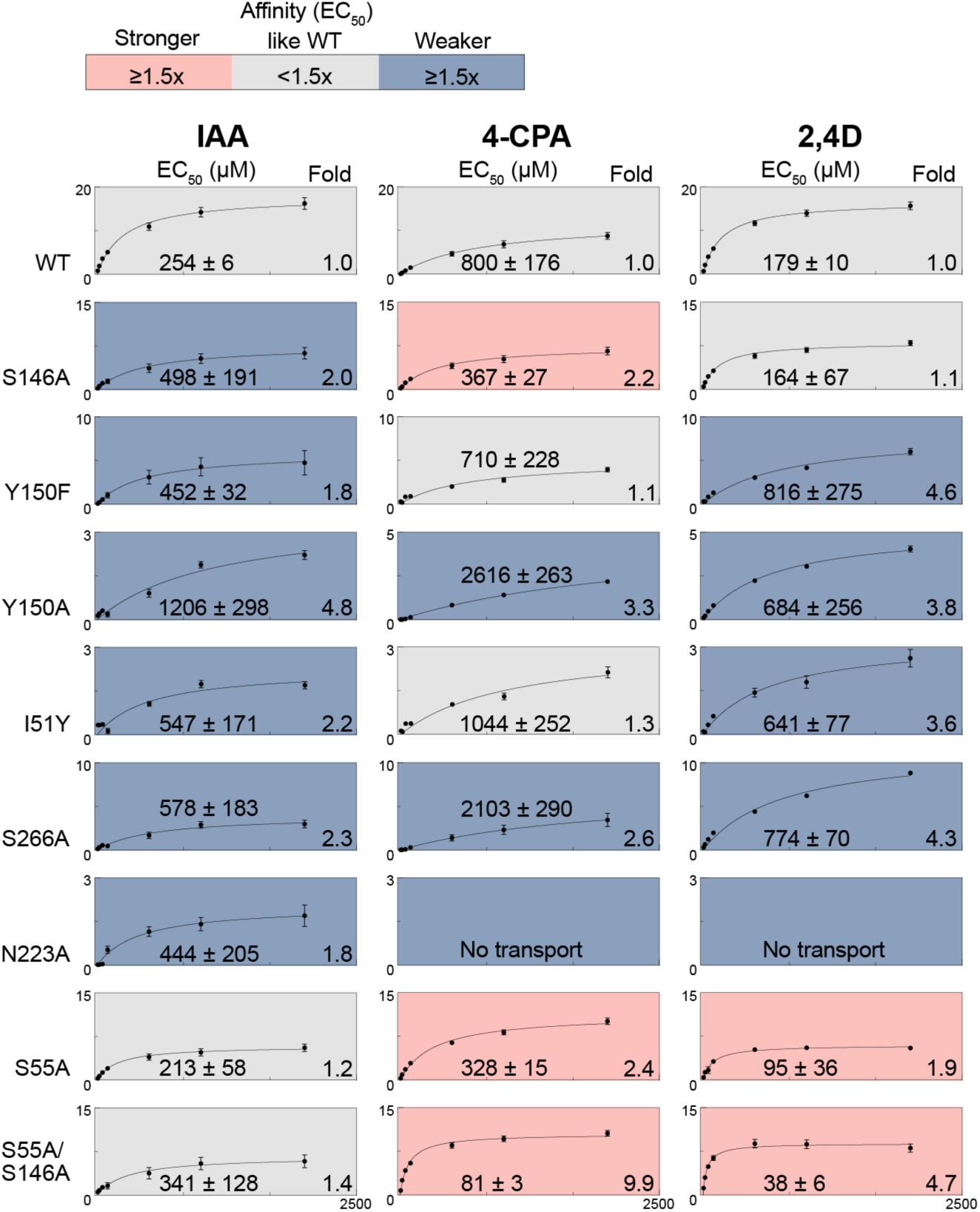
Identification of key residues required for substrate binding. A) *EC_50_* of peak current response as a function of the substrates indicated derived from the full measurements. A Michaelis–Menten model is fit to describe kinetics (Extended Data Table 4) Red color indicates ≥ 1.5x higher affinity than WT; blue colour ≥ 1.5x lower affinity than WT; grey color < 1,5x change from WT affinity. No transport means that a Michaelis-Menten-Model could not be fit. Indole-3-acetic acid (IAA); 2,4-Dichlorphenoxyacetic acid (2,4D); 4-Chlorphenoxyacetic acid (4-CPA).

The structures suggest that Asn223 is a critical residue that delineates the vestibule from the binding pocket in the inward conformation. In contrast to previous work on PIN1 ^17^, we find that the N223A mutant reduces affinity for IAA only slightly, but completely abolishes binding of 2,4-D and 4-CPA. This result suggests that once the substrate has flipped from the prebinding position into the binding pocket, Asn223 is required to keep the substrate in place (Extended Data Fig. 5E). In the outward open substrate-bound state Ser55 is near the 4-Cl group (Fig. 3B and E). We reasoned that mutating Ser55 to an Alanine would accommodate a better fit of 2,4-D and 4-CPA as well as support the more hydrophobic nature of the halogenated phenyl ring. Indeed, the S55A mutant show higher affinity (1.9- to 2.4-fold) for the herbicides, whereas the affinity for IAA remains unchanged. To further support the notion that increased hydrophobicity and size of the binding pocket size can selectively improve binding of the chlorinated auxin herbicides, we tested the double mutant S55A/S146A (Fig. 4), which increase affinity towards chlorinated auxin herbicides even further (4.7- to 9.9-fold), again with no effect on affinity of the natural auxin IAA. Taken together the results suggest that by combining chemical variability in substrates and protein mutagenesis it will be possible to design novel auxins and engineer resistance to these compounds by a structure guided approach.

## DISCUSSION

We have shown that synthetic auxins are transported by PIN proteins both *in planta* and *in vitro*, and have explored the biochemical, structural, and chemical parameters that govern transport. Specifically, we have used cryo-EM to describe changes in substrate binding as the protein undergoes conformational changes associated with transport. We have used structure-guided mutagenesis to evaluate principal components of the binding pocket that recognize and interact with substrates and have used chemical biology to elucidate substrate determinants of binding affinity. Our work highlights key chemical and structural elements of both herbicides and auxin transporters that can be adjusted to augment or prevent herbicide distribution in plants.

Structural analysis of transport under turnover conditions has been demonstrated for primary active transporters (e.g., ^23,24^). This has been facilitated by their relatively slow turnover rate and by the ability to use ATP to initiate the transport cycle prior to flash freezing of cryo-EM samples. While turnover conditions cannot as easily be established for uniporters, we demonstrate an analogous approach here for PIN8, where a single sample can contain a multitude of different conformational states that represent a number of transport cycle intermediates.. In the case of 4-CPA, the structures provide a comprehensive mapping of molecular interactions between a PIN transporter and an auxin herbicide as it transits across the membrane. The structural data show that transport is a multi-step process with auxin first recognized in the vestibule before nestling into the binding pocket. Occupation of this pocket induces the change to the outward state, where the binding steps are reversed. An occluded state with substrate trapped in the binding pocket has not been observed, suggesting that it is a short-lived, unstable state, at least in the presence of 4-CPA.

Binding of phenoxyacetic acid herbicides to PINs is very similar to the natural substrate, IAA (Extended Data Fig. 7). The substrates enter the vestibule, and the carboxylate is coordinated directly by Asn117 and indirectly by Tyr150, Ser146 and Gln145 via a water molecule. These findings are consistent with previous work in which we determined Asn117 to be critical for the coordination of IAA’s carboxylate and for transport activity in PIN8 ^16^. In the current work, 2,4-D is seen in a prebinding position with its carboxyl group interacting with Asn117 and with crossover dipoles from the nearby half-helices neutralizing its negative charge; like IAA, 2,4-D is transported with an *EC_50_* ∼200 μM and the N117A mutation abolishes function (Fig. 1A, Fig. 4 and Extended Data Fig. 5D). Although this general location of 2,4-D is similar to the IAA position in the previously published structure of PIN1 the pose is different and closer to that of IAA in the previously published structure of PIN3 ^14,15,17^. Based on SSM analysis, the 4-Cl group and its interaction with Asn223 is critical. We show that Ser55 and Tyr150 play the critical role in the binding pocket. Removing the polar group of Ser55 accommodates a better fit of the halogenated substrate while the non-halogenated ring of IAA is accommodated with no changes in affinity. Addition of Cl-groups to the benzyl ring reduces the electron density of the π-electrons over the benzyl ring ^25^ until a π-configuration similar to the indole ring is achieved. Notably this configuration is not stabilized by methyl groups in the ring (Fig. 2A).

Like many other transporters employing the elevator mechanism, PINs assemble into dimers, and this assembly has been hypothesized to provide potential for cooperative action during transport. Here we show that, at least for PIN-FORMED auxin transporters, the conformation of protomers within the dimeric assembly are uncorrelated. This observation is inconsistent with cooperativity; rather, it suggests that each protomer operates independently. Regarding the elevator mechanism, superposition of transport and scaffold domains of all the structures show that PIN transport involves rigid body movements, without local changes in these individual domains (Fig. 1D and 3A-C, Extended Data Fig. 8); transition from inward to outward states involves a translation of the transport domain in a direction normal to the membrane plane while the scaffold domain remains fixed within this plane and anchored to its dimeric partner molecule.

In conclusion, we have shown that phenoxyacetic acid auxin herbicides are transported by PINs. We have used cryo-EM to study PIN8 and have captured five key states during transport of the herbicide 4-CPA. This structural analysis elucidates the steps of substrate binding and release, provides a model to explain how chemical diversity of synthetic auxins can be accommodated, and indicates that dimeric transporters with a crossover fold can operate asymmetrically with an apparent lack of cooperativity. Future research will show whether other auxin herbicides are also PIN substrates and if the determinants of substrate affinity can be generalized to include other halogenated auxin herbicides such as Dicamba or Picloram ^2^. In this way, our elucidation of the determinants of substrate affinity may be used to identify novel environmentally safer auxin herbicides with higher systemic mobility for sustainable agriculture.

## Acknowledgements

We acknowledge the EMBION Cryo-EM Facility at iNANO, Aarhus University (application #0137), where data was collected with the assistance of Andreas Bøggild, Taner Drace and Thomas Boesen. We also thank eBIC for Cryo-EM data collection (application BI21404). We acknowledge access to the computational infrastructure at the Center for Structural Biology. U.Z.H. is funded by the Deutsche Forschungsgemeinschaft (HA3468/6-3). KWS SAAT SE & Co is acknowledged for financial support to UZH. S.K.A. and B.P.P. are supported by funding from the Technical University of Munich – Institute for Advanced Study. This project has received funding from the European Research Council (ERC) under the European Union’s Horizon 2020 research and innovation programme (grant agreement No 101000936) to B.P.P. D.L.S. is supported by the National Institutes of Health (Grant Agreement No. R35 GM144109).

## Author Contributions

Activity assays: LS, SKA, UZH

Sample preparation: KLU

EM data: KLU, DLS, BPP

Data analysis: LS, KLU, SKA, DLS, BPP, UZH

Manuscript preparation: KLU, LS, DLS, BPP, UZH

## Data Availability

Materials are available upon request. Atomic models have been deposited in the Protein Data Bank (PDB) and Cryo-EM maps have been deposited in the Electron Microscopy Data Bank (EMDB).

2,4-D inward prebinding (vestibule): PDB 9G0W and EMD-50950.

4-CPA inward prebinding (vestibule) (I_4CPA prebinding_/O_empty_): PDB 9G0X and EMD-50951.

4-CPA outward binding (O_4CPA binding_/O_empty_): PDB 9G0Z and EMD-50952.

4-CPA outward partly released (O_4CPA, partlyreleased_/I_empty_): PDB 9G10 and EMD-50953.

## Supplementary Materials

Figs. S1 to S8

Table S1 to S4

## Additional information

Correspondence and requests for materials should be addressed to B.P.P. or U.Z.H.

## Materials and Methods

### Solid supported membrane-based electrophysiology assays

SSM, using a SURFE2R N1 from Nanion Technologies was conducted as described as described ^16^. In brief, soy polar lipid mix (38% phosphatidyl choline, 30% phosphatidyl ethanolamine, 18% phosphatidyl inositol, 7% phosphatidic acid and 7% other soy lipids) and 1-palmitoyl-2-oleoyl-sn-glycero-3-phosphocholine (POPC) were purchased from Avanti. Liposomes were prepared in Ringer solution without Ca^2+^ (115 mM NaCl, 2.5 mM KCl, 1 mM NaHCO_3_, 10 mM HEPES pH7.4, 1 mM MgCl_2_) and homogenized using a Lipsofast (Avestin Inc) with a 400 nM pore size. Triton X-100 was added to the liposomes to a final concentration of 1% (v/v). Protein was added to liposomes to a calculated liposome:protein ratio (LPR) of 10 unless stated otherwise. The detergent was removed using 400 mg/mL Bio Beads (BioRad) overnight at 4°C in a rotary shaker. Proteoliposomes were frozen in liquid nitrogen and kept at -80°C until use. Proteoliposomes were diluted 1:5 in Ringer solution without Ca^2+^, sonicated five times and then applied to the sensors by centrifugation (30 min, 3,000 g, 4°C). Non-activating buffer was Ringer solution without Ca^2+^ as described unless specified otherwise and activating buffer contained the substrate of interest. In most instances we used a single solution exchange experiment. In this case proteoliposomes, immobilized on the supported membrane are kept in non-activating buffer as specified. At the beginning of the experiment non-activating buffer is exchanged for fresh identical non-activating buffer and after 1 sec activating buffer (same buffer containing substrate) is added. After another sec buffer is exchanged to non-activating buffer again. Current response is recorded throughout the entire 3 sec. For competition or inhibition, the respective compound was present in non-activating and activating solution. Each experiment was performed on at least two individual sensors. On each sensor each measurement consists of three technical replicates where the mean is calculated. *K_D_* values were determined as described previously ^16^. Briefly, the binding current induced by 100 µM NPA was recorded in presence of different concentrations of the substrate under investigation ranging between 0 µM and 1000 µM present in the activating solution and the non-activating solution (Extended Data Fig. 5B). Representative traces are shown for 0, 50 and 1000 µM 2,4-D (Extended Data Fig. 5B). The maximum peak current was then plotted as a function of the concentration of the substrate under investigation and plotted against the degree of inhibition and fitted using a one-site binding model using GraphPad Prism V10.2. The half maximal concentration corresponds to the *K_D_* of the substrate under investigation.

### Oocyte efflux assays

Oocyte efflux experiments were carried out as described ^16,21,26^. Briefly, oocytes were injected with 150 ng transporter cRNA without or with 75 ng kinase cRNA. [^3^H]-IAA (25 Ci/mmol) was purchased from RC Tritec. [^3^H]-2,4-D (25 Ci/mmol) was purchased from ARC, St. Louis, MO. Oocytes were injected with substrate to reach an internal concentration of 1 µM, corresponding to 100%. Residual radioactivity was determined for each individual oocyte by liquid scintillation counting after the time points indicated and are expressed relative to the initial 100%. Each time point represents the mean and SE of ten oocytes. To calculate the relative transport rate in % per minute linear regression was performed. Each data point in Fig.s 1B and C, and Extended Data Fig. 2B and E represents the transport rate of one biological replicate using oocytes harvested from different Xenopus females.

### Stem assays

Arabidopsis stem transport assays were carried out as described ^21^. Briefly, 2 cm stem sections were cut above the rosette of 5-week-old plants and placed in inverted orientation into auxin transport buffer containing 500 pM IAA, 1% (wt/vol) sucrose, 5 mM 2-(N-morpholino)ethanesulfonic acid (MES), pH 5.5 with or without 100 μM 1-N-naphthylphthalamic acid (NPA). After two hours the stems were transferred into auxin transport buffer containing 417 nM [^3^H]-IAA or [^3^H]-2,4-D as a substrate in the absence and presence of NPA. After two hours, 5 mm segments were dissected, and the substrate was quantified using a liquid scintillation counter.

### Cryo-EM sample preparation

The protocol for expression and purification of AtPIN8 protein (Uniprot: Q9LFP6) in LMNG was described previously ^16^. All point mutants were generated using Q5 Site-Directed mutagenesis kit (New England Biolabs). The C-flat Holey Carbon grids (CF-1.2/1.3, Cu-300 mesh) were glow-discharged for 45 s at 15 mA in a GloQube Plus (Quorum). The 2,4-D sample was prepared by mixing freshly purified PIN8 at 9.4 mg/mL in G-Buffer (50 mM Tris pH 7.5, 0.15 M NaCl, 0.0006% LMNG) with 2,4-D (dissolved in DMSO) to reach a ∼15 mM final concentration. After 1 h of incubation on ice, the sample was centrifuged at 16000 g for 15 minutes prior to blotting to remove precipitation. A drop of 4 µl of each sample was applied to the glow-discharged grids, blotted with a Vitrobot Mark IV (ThermoFisher Scientific) using the following settings (temperature 4oC, 100% humidity, blot time of 4s, blot force - 1) and vitrified in liquid ethane. For 4-CPA, after mixing protein (8 mg.ml-1) with 25 mM 4-CPA (dissolved in DMSO) on ice, sample was rapidly applied on the grid without centrifugation. We did not accurately estimate the incubation time during blotting but the incubation window for the collected grid is between 10 to 20 minutes.

### Image collection and data processing

A Titan Krios G3i microscope (ThermoFisher Scientific) operating at 300 kV and equipped with a BioQuantum Imaging Filter (energy slit width of 20 eV) with a K3 detector (Gatan) was used to collect the movies. The datasets were acquired using automated acquisition EPU at nominal 130,000 magnification corresponding to a physical pixel size 0.647 Å. For all datasets, the movies were saved in super-resolution pixel size and binned 2x in EPU back to the nominal pixel size.

Gain normalize micrographs were imported into cryoSPARC and processed in for patch motion correction, patch contrast transfer function (CTF) estimation and particle picking with blob picker ^27^. After several rounds of particle cleaning, an initial preliminary volume map was used to create templates for template picking.

From a full data set of 2,4-D PIN8 with 8,473 movies, template-picking provided a total of 3,173,516 particles. The full stack of particles was divided into four smaller particles stack to reduce computational time. After 2 rounds of 2D classifications, manually selected 2D classes from each stack were combined, yielded a total of 875,481 particles, and re-extracted in a box size of 416 pixel (Fourier-crop to box size 208) and divided into two particles stack.

Here, we design an ENRICH-workflow to recover a maximum number of good particles that have been lost during particles cleaning jobs. The initial particles stack will be processed independently three times and recombined after 2 rounds of 2D classifications, providing a total of 662,418 particles, served as input for ab-initio model reconstruction. Of four ab-initio models, the best representative *ab-initio* was served as “good” template volume. The “bad” template volume was generated by low pass filtering the “good” template volume to 40Å. The corresponding particles stack from good *ab-initio* class (209,920 particles) were sorted with rounds of heterogeneous refinement using 3 “good” and 3 “bad” template volumes and processes independently 4 times. The good particle stack from each individual heterogenous refinement were recombined, resulting in a total of 281,146 particles. Following an *ab-initio* model reconstruction, particles from good class were re-extracted and were used for non-uniform refinement with C1 symmetry imposed and resulted in a global 3.5 Å resolution map.

The entire 4-CPA PIN8 dataset comprised of 14,655 movies and template-picking yielded a total of 8,093,919 particles. The full stack of particles was divided into four smaller subsets. After 2D classification, selected 2D classes particles were combined, resulting in a total of 1,564,727 particles. After two rounds of *ab-initio*, two classes with clear transmembrane helices were selected for further processing. Preliminary model fitting into reconstructed *ab-initio* volume suggested the presence of an asymmetric (class 0) and symmetric (class 2) configuration of each monomer in the PIN dimer.

Particles belonging to *ab-initio* class in which each monomer adopted an outward-inward configuration was further processes by a second *ab-initio* reconstruction and focus-3D classification with a spherical mask (15Å, dilation radius 2 Å, soft padding width 5) centring around the outward binding pocket in the outward monomer. Particles from each class were further used for NU-refinement with C1 symmetry imposed and only one class resulted in a global 3.4 Å resolution map with well-defined ligands density in the outward binding pocket (Map 1).

To avoid the recycling of particles belonging to the first map, the initial particles stack from the main *ab-initio* was subtracted with particles belonging to map 1. The remaining stack (992,336 particles) were subjected to heterogenous refinement job using volume from the previous main *ab-initio* as template. Similarly, particles belonging to class in which each monomer adopted an outward-outward configuration were processed using a focus-3D classification with a spherical mask centring around the outward binding pocket in one outward monomer. Particles from each class were further used for NU-refinement with C1 symmetry imposed and only one class resulted in a global 3.3 Å resolution map with well-defined ligands density in the outward binding pocket (Map 2). The same strategy was used with a focus mask centring around the inward binding pocket. Isolated particles stack is further refined to 3.4 Å resolution map (C1 symmetry) with well-defined ligand density in the inward binding pocket (Map 3).

### Model building and refinement

The published apo-PIN8 structure (PDB:7QP9) served as initial model for docking into the map for 2,4-D and 4-CPA using Chimera ^28^. Molecular Dynamics based geometry fitting using MDFF (Trabuco et al., 2008) was carried out by Namdinator ^29^ After this, models were manually adjusted by iterative model building in Coot combined with real space refinement using Phenix, initially with an Amber force-field molecular dynamic refinement ^30^. The coordination of lipids and the ligand 2,4-D, 4-CPA and DMSO was obtained from GRADE (https://grade.globalphasing.org/). Structure geometry was validated by monitoring MolProbity, CaBLAM and Ramachandran-Z analysis (Rama-Z) ^31–33^. Statistic of the model refinement can be found in Table 1Fig.s were prepared using PyMOL Molecular Graphics System (Schrödinger, LLC) and Chimera ^28^.

### Statistical analyses

Statistical analyses were performed using GraphPad Prism 10.2. Parameters of the individual tests and results are shown Extended Data Tables 1 to 3.

---

**Extended Data Fig. 1:**
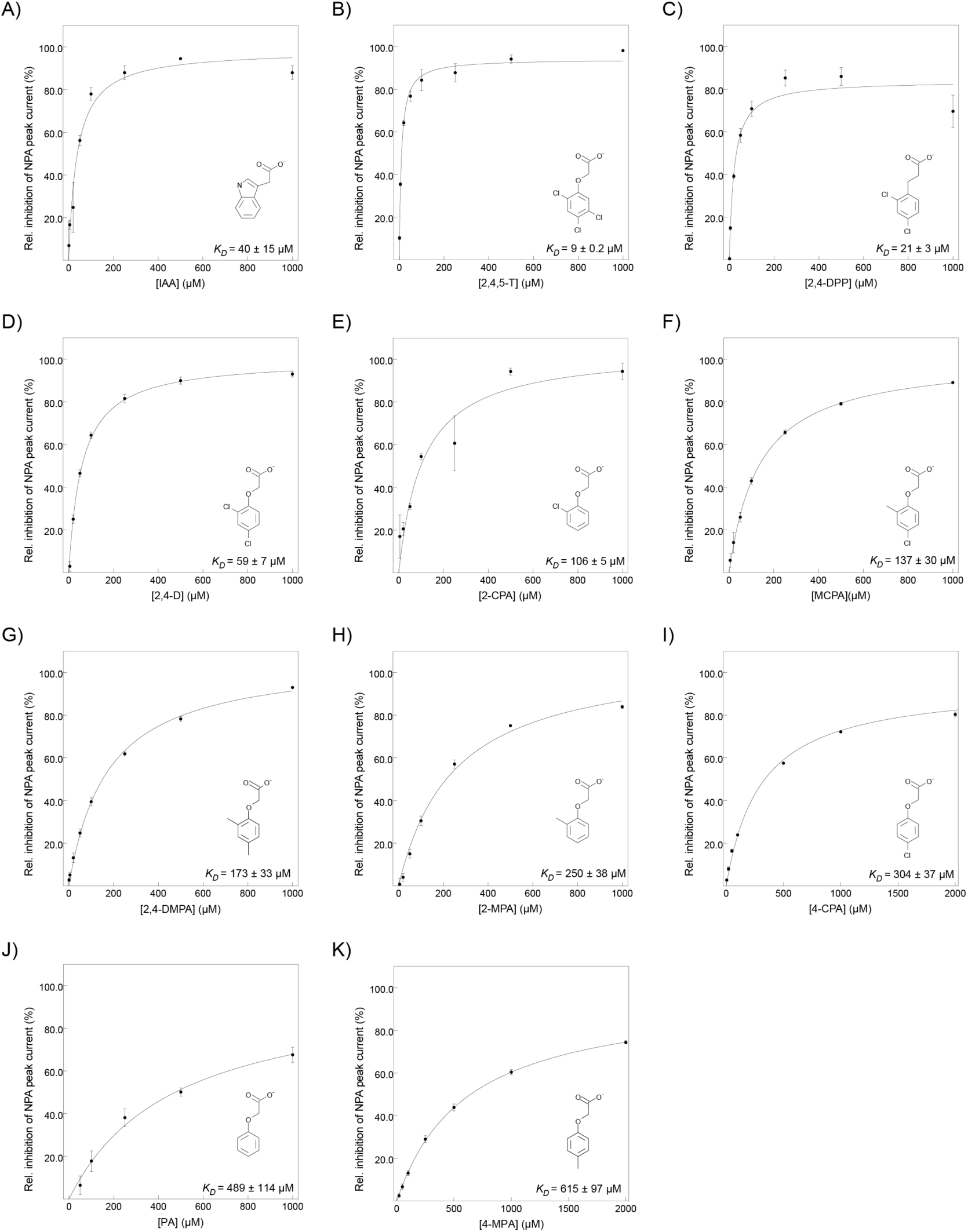
Full *K_D_* measurements of substrates. *K_D_* of substrate binding by PIN8 determined by SSM electrophysiology. Relative inhibition of Peak current response elicited by 100 µM NPA by [substrate] indicated. A Michaelis–Menten model is fit to describe kinetics; data points represent mean ± s.e.m. A) IAA (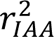 = 0.91, 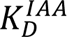 40 ± 15 μM; n = 3 individual sensors) B) 4-CPA (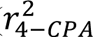 = 0.99, 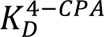 304 ± 37 μM; n = 3 individual sensors) C) (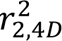 = 0.99, 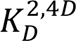 59 ± 7 μM; n = 3 individual sensors) D) 2,4,5-T (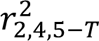 = 0.98, 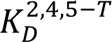 9 ± 0.2 μM; n = 3 individual sensors) E) MCPA (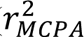 = 0.99, 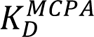 137 ± 30 μM; n = 3 individual sensors) F) PA 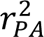 = 0.95, 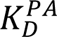 489 ± 114 μM; n = 3 individual sensors) G) 2-CPA (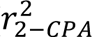 = 0.89, 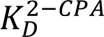 106 ± 5 μM; n = 3 individual sensors) H) 2-MPA (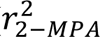 = 0.99, 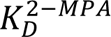 250 ± 38 μM; n = 3 individual sensors) I) 2,4-DPP (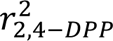 = 0.93, 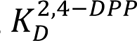 21 ± 3 μM; n = 3 individual sensors).

**Extended Data Fig. 2:**
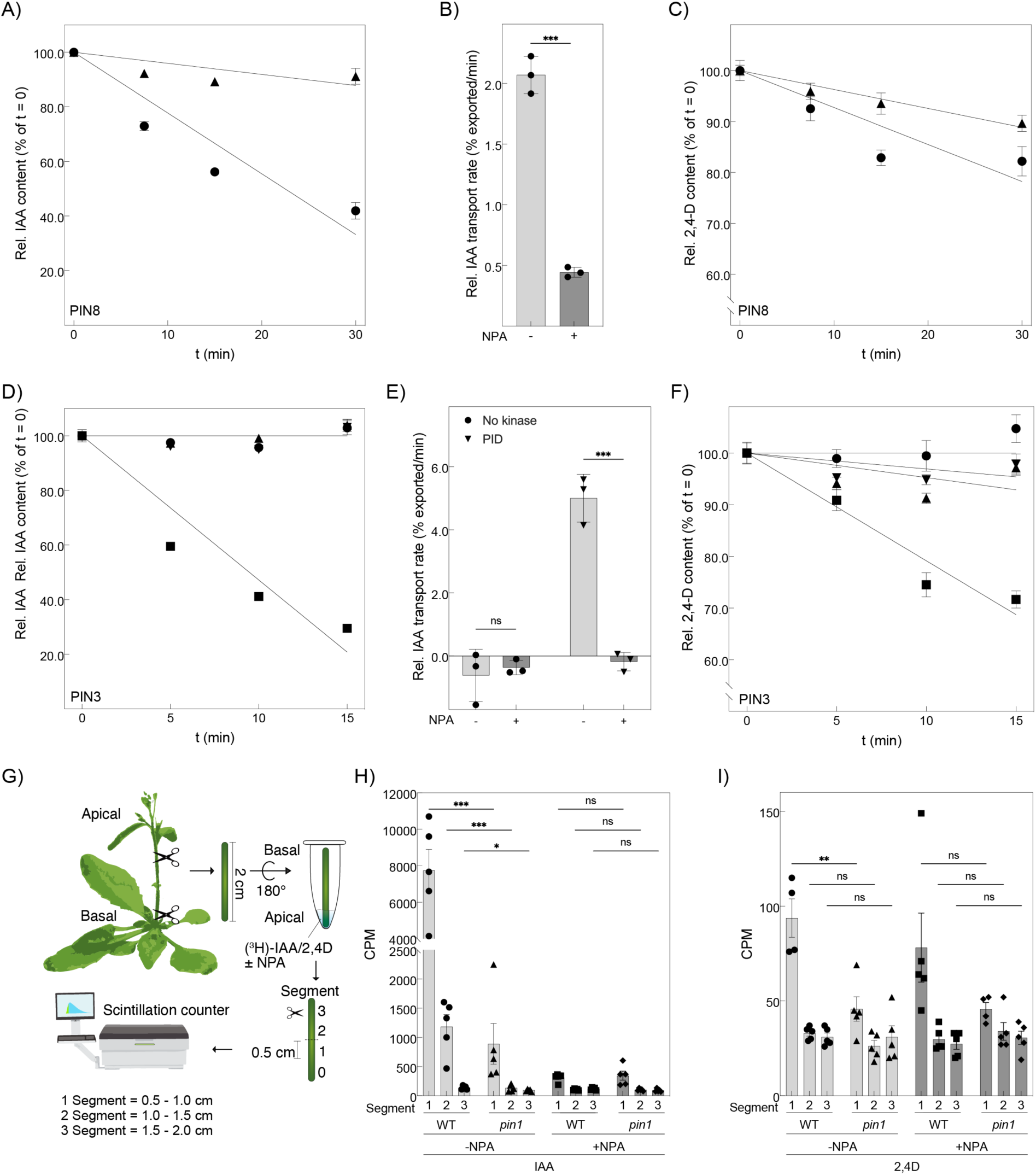
PIN mediated IAA and 2,4D export from Xenopus oocytes and basipetal auxin transport in inflorescence Arabidopsis stems. A) Time course of [3H]-IAA export by PIN8 from oocytes. Data of a typical experiment. Relative IAA content of oocytes expressing PIN8 in the presence (▴) or absence (•) of 10 μM NPA internally determined at the time indicated after substrate injection. Initial internal IAA concentration was 1 μM. n = 10 oocytes at each time point. Data points are mean ± SE. B) Relative transport rates of [3H]-IAA export by PIN8 in presence (dark grey) or absence (light grey) of 10 μM NPA as indicted from oocytes calculated from three independent time course experiments. Data points are individual experiments and bars represent mean ± SD. Groups were compared by a two-sided unpaired Student’s t-test; P < 0.0001. C) Time course of [3H]-2,4D export by PIN8 from oocytes. Data of a typical experiment. Relative IAA content of oocytes expressing PIN8 in the presence (▄) or absence (•) of 10 μM NPA internally determined at the time indicated after substrate injection. Initial internal 2,4D concentration was 1 μM. n = 10 oocytes at each time point. Data points are mean ± SE. D) Time course of [3H]-IAA export by PIN3 from oocytes. Data of a typical experiment. Relative IAA content of oocytes expressing PIN3 without kinase in the presence (▄) or absence (•) of 10 μM NPA internally and PID in the presence (▾) or absence (▴) of 10 μM NPA internally determined at the time indicated after substrate injection. Initial internal IAA concentration was 1 μM. n = 10 oocytes at each time point. Data points are mean ± SE. E) Relative transport rates of [3H]-IAA export by PIN3 with or without PID in presence (dark grey) or absence (light grey) of 10 μM NPA as indicted from oocytes calculated from three independent time course experiments. Data points are individual experiments and bars represent mean ± SD. Groups were compared by a two-sided unpaired Student’s t-test. n.s. not significant (P =0.99), *** P < 0.0001. F) Time course of [3H]-2,4D export by PIN3 from oocytes. Data of a typical experiment. Relative 2,4D content of oocytes expressing PIN3 without kinase in the presence (▄) or absence (•) of 10 μM NPA internally and PID in the presence (▾) or absence (▴) of 10 μM NPA internally determined at the time indicated after substrate injection. Initial internal 2,4D concentration was 1 μM. n = 10 oocytes at each time point. Data points are mean ± SE. G) Illustration of the stem transport assay measured in inflorescence stems of 5-week-old Arabidopsis plants. 2 cm stem sections were cut above the rosette of 5-week-old plants and placed in an inverted orientation into auxin transport buffer containing [3H]-IAA or [3H]-2,4D as a substrate in the absence and presence of NPA. After two hours, 5 mm segments were dissected, and the substrate was quantified using a liquid scintillation counter. Segment 1 represents 0.5 to 1 cm, Segment 2 represents 1 to 1.5 cm and Segment 3 represents 1.5 to 2 cm of the stem, from apical to basal position. H) [3H]-IAA content in segments of Col-0 wildtype and pin1 mutant plants in the absence and presence of NPA as indicated. Data points are segments of individual plant stems and bars represent mean ± SD. Groups were compared by a two-sided unpaired Student’s t-test. For individual p-values see Extended Data Table 2.I) [3H]-2,4-D content in segments of Col-0 wildtype and pin1 mutant plants in the absence and presence of NPA as indicated. Data points are segments of individual plant stems and bars represent mean ± SD. Groups were compared by a two-sided unpaired Student’s t-test. For individual p-values see Extended Data Table 2.

**Extended Data Fig. 3:**
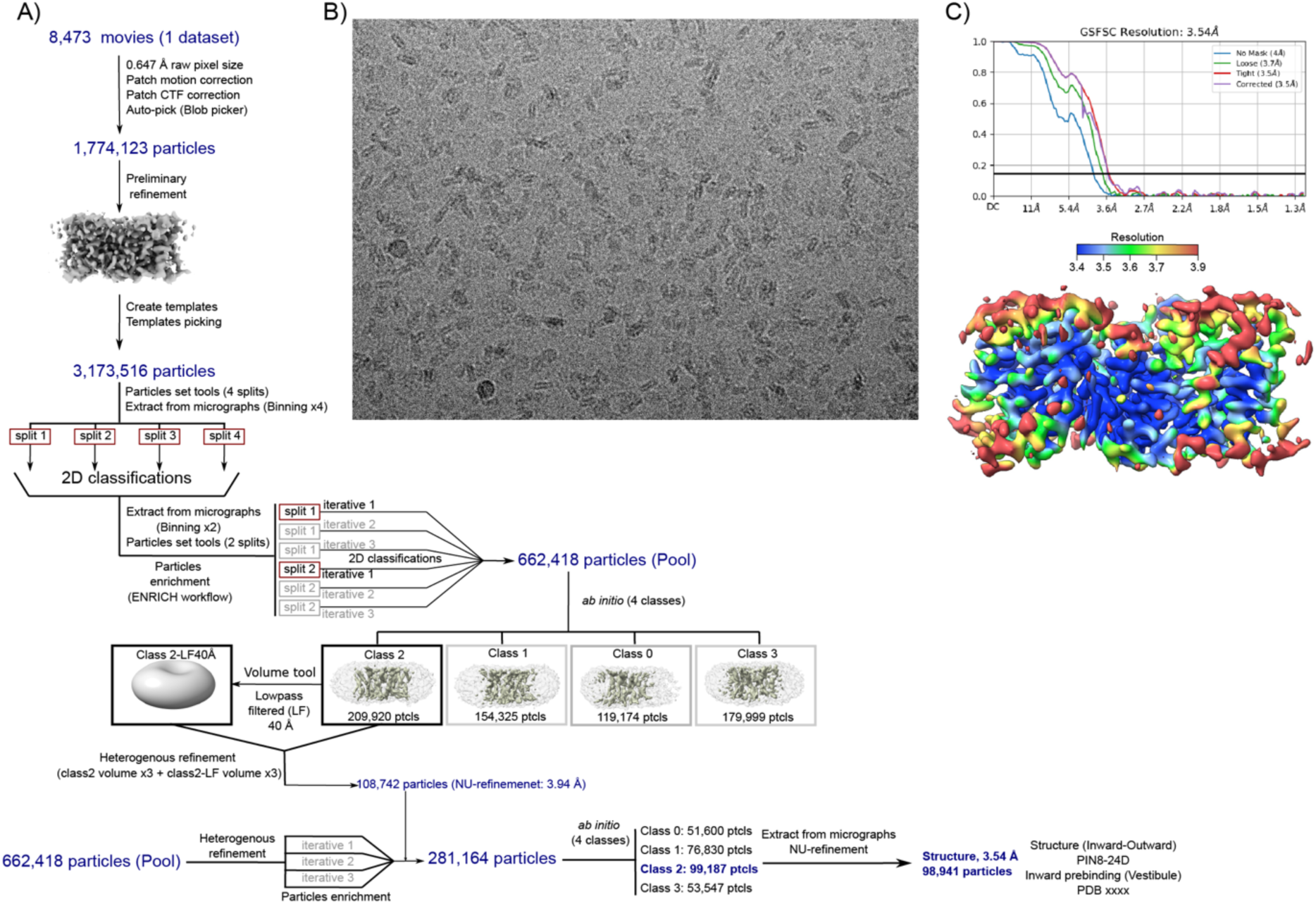
Image processing and reconstruction for 2,4-D bound PIN8. Workflow of image processing and 3D reconstruction in cryoSPARC, including a motion corrected micrograph from Titan Krios microscope using a K3 detector, sharpened density map from the final non-linear refinement coloured by local resolution. Corrected curve of the global Fourier shell correlation (FSC) indicates 3.54 Å based on the 0.143 gold-standard criterion.

**Extended Data Fig. 4:**
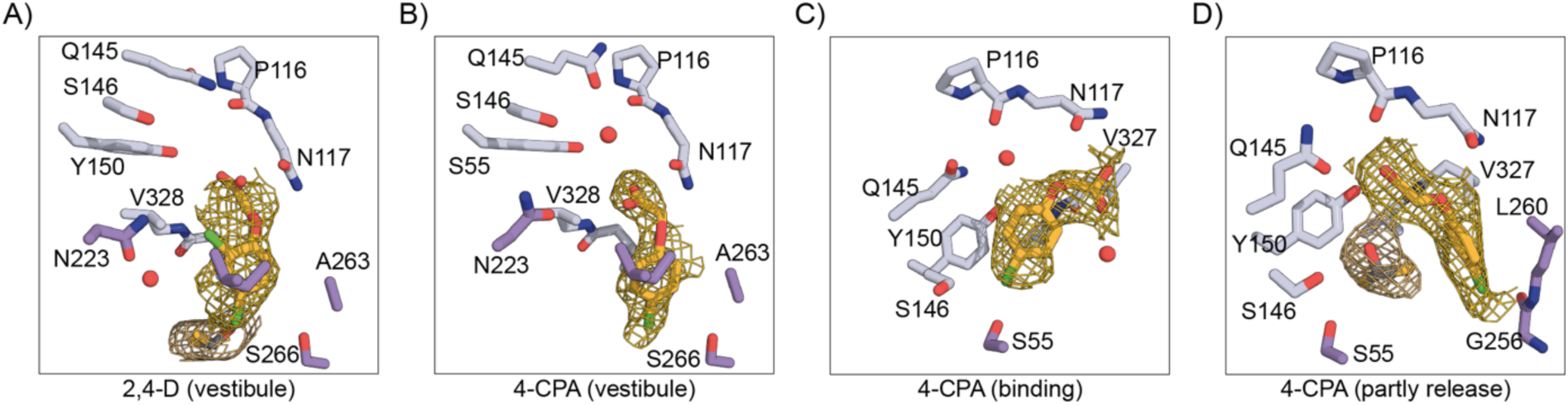
Structures of PIN8 with bound 2,4-D or 4-CPA. Close-up view of 2,4-D and 4-CPA map density and the residues interacting with the compound.

**Extended Data Fig. 5:**
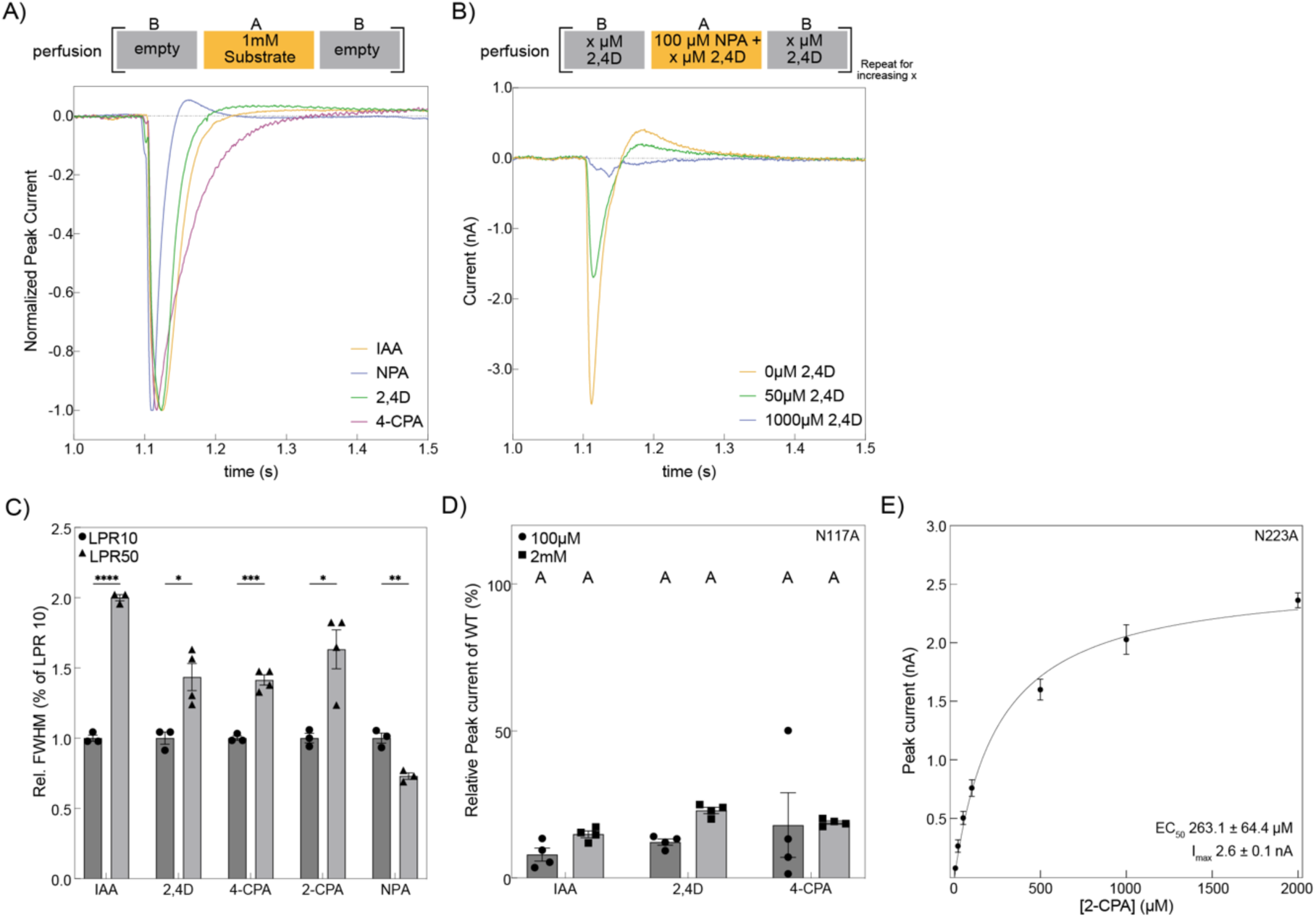
Transport of phenoxyacetic acids and identification of critical residues for Substrate interaction. A) Normalized representative current traces elicited by 1mM of the substrates. The perfusion protocol is shown on the top. B) Raw current traces of three representative solution exchange experiments used for *K_D_* determination of 2,4D. For clarity the graph only shows three of the concentrations used to obtain the *K_D_*. The perfusion protocol is shown on the top. C) Full width half maximum (FWHM) from the Peak current response determined by SSM electrophysiology on the wild-type PIN8 proteoliposomes of LPR10 (•) and LPR50 (▴) derived from 2mM of indicated substrate. Data are mean ± s.e.m. of n = 3 individual sensors. Groups were compared by a two-sided unpaired Student’s t-test (IAA_LPR10_ versus IAA_LPR50_: P < 0.0001; 2,4D_LPR10_ versus 2,4D_LPR50_: P = 0.0148; 4-CPA_LPR10_ versus 4-CPA_LPR50_: P = 0.0003; 2-CPA_LPR10_ versus 2-CPA_LPR50_: P = 0.0127; NPA_LPR10_ versus NPA_LPR50_: P = 0.0032). D) Relative Peak current response compared to WT elicited by 100μM (•) or 2mM (▄) of indicated substrates using SSM electrophysiology on PIN8 N117A proteoliposomes. Data are mean ± s.e.m. of n = 3 individual sensors. There are no significant differences between the substrates at both concentrations used (One way ANOVA p = 0.3). Letters indicate the same significance level. E) Peak current response as a function of [2-CPA] determined by SSM electrophysiology on PIN8 N223A proteoliposomes. A Michaelis–Menten model is fit to describe kinetics (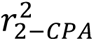= 0.98; 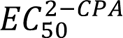 263 ± 64 μM; 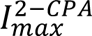 3 ± 0.1 nA; data points represent mean ± s.e.m.; n = 3 individual sensors).

**Extended Data Fig. 6:**
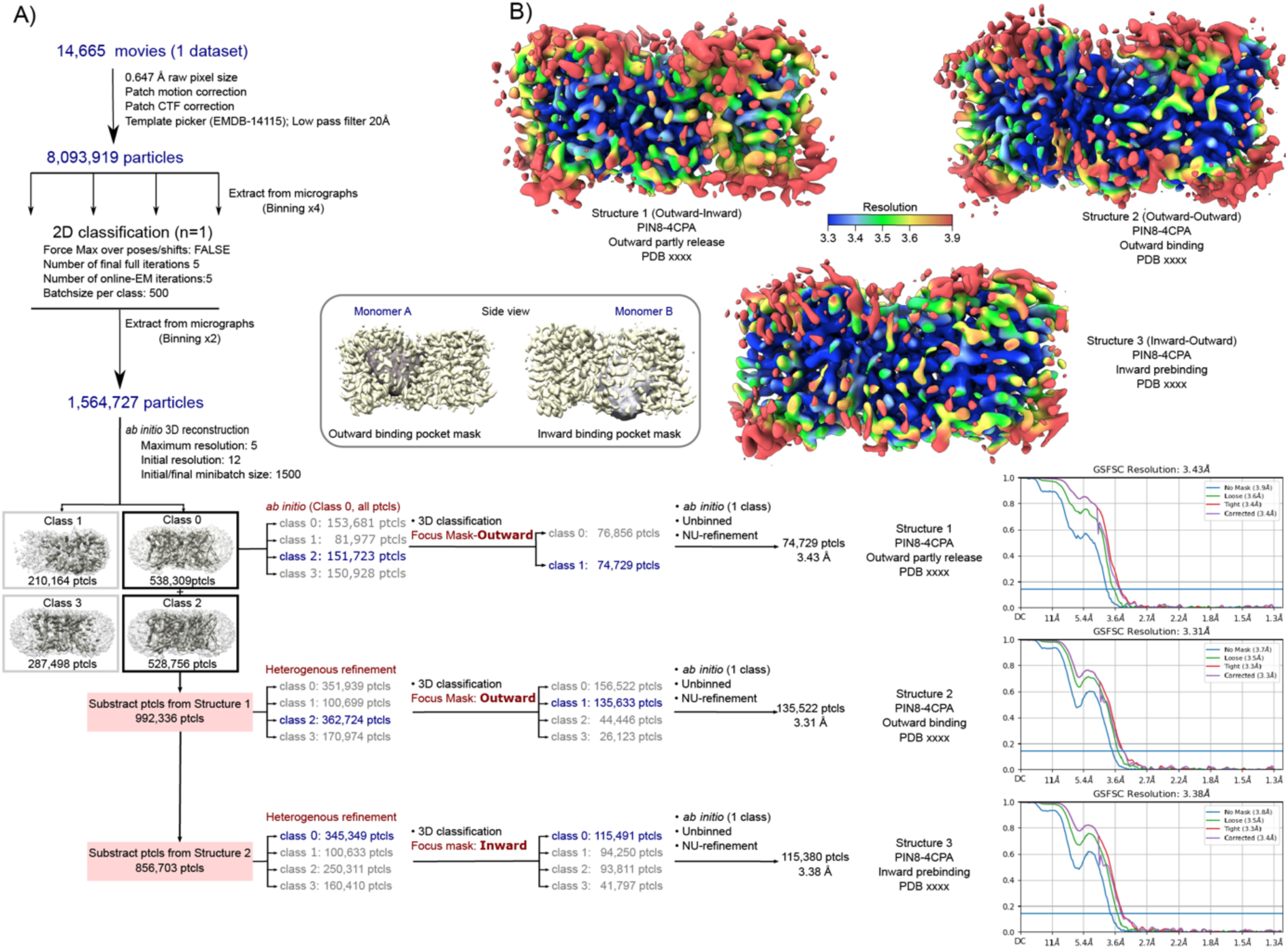
Image processing and reconstruction for 4-CPA bound PIN8. Workflow of image processing and 3D reconstruction in cryoSPARC, including a motion corrected micrograph from Titan Krios microscope using a K3 detector. B) Sharpened density map from the final non-linear refinement colored by local resolution. Corrected curve of the global Fourier shell correlation (FSC) indicates 3.43 Å (Structure 1), 3.31 Å (Structure 2), 3.38 Å (Structure 3) based on the 0.143 gold-standard criterion. The cryo-EM experiment with this sample was repeated 5 times with data collection 1 time.

**Extended Data Fig. 7:**
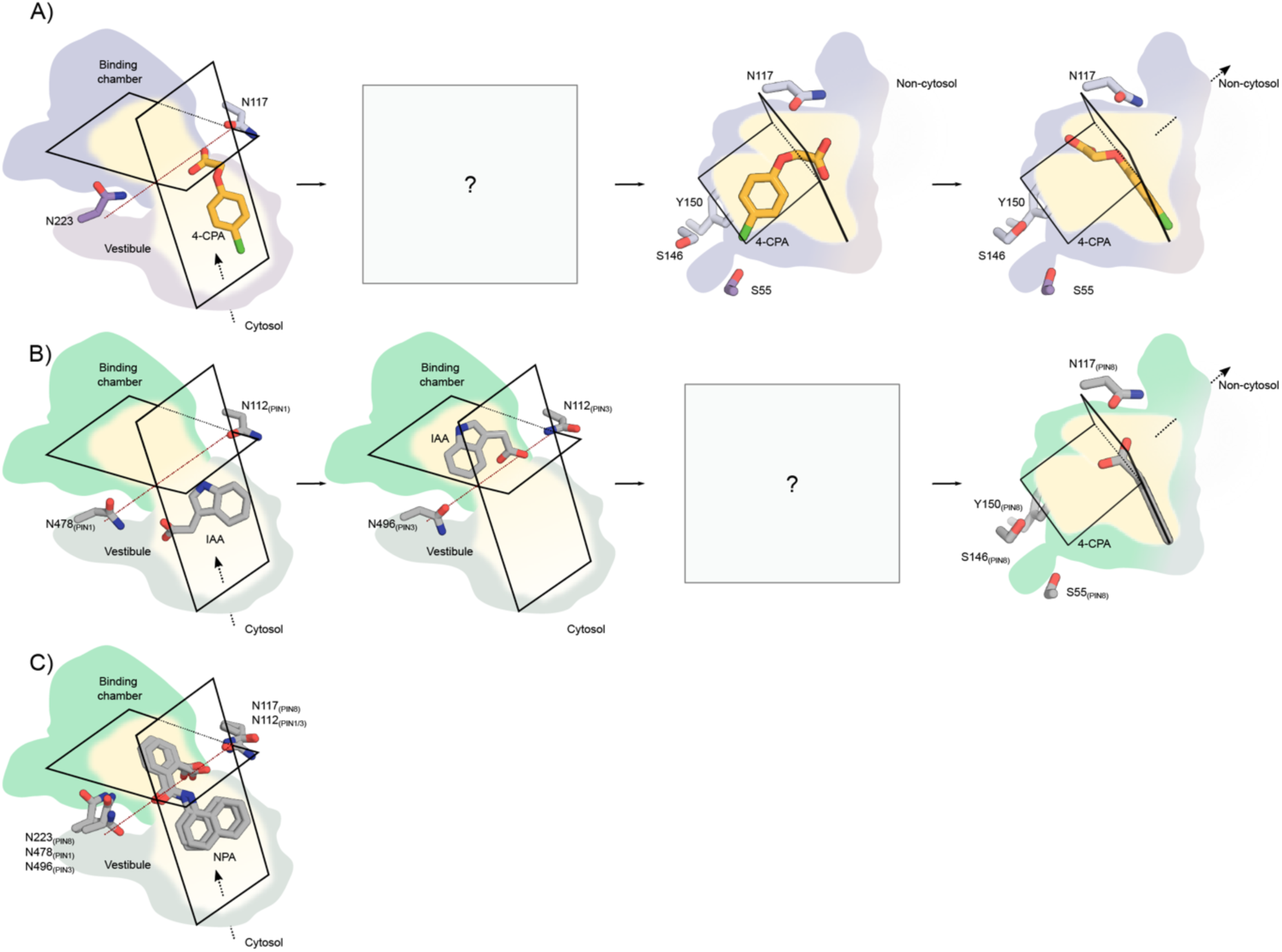
Substrate binding and interaction. Structures are superposed on the scaffold domain. The inward-facing pocket is divided into the binding chamber and a vestibule. The red broken line running from Asn117 to Asn223 defines the boundary between vestibule to binding chamber. (A) 4-CPA binding position and poses in the prebinding in the PIN8-inward conformation, binding and partly release in the PIN8-outward conformation. Square with question mark represents the missing state. (B) IAA binding position and poses in the prebinding in the PIN1-inward conformation, binding in the PIN3-inward conformation, and partly release in the PIN8-outward conformation. Square with question mark represents the missing state. (C) Overlay of NPA binding poses from PIN8, PIN1, PIN3. NPA binds in both the binding chamber and the vestibule and forms hydrogen bonds to both Asn117 and Asn223.

**Extended Data Fig. 8:**
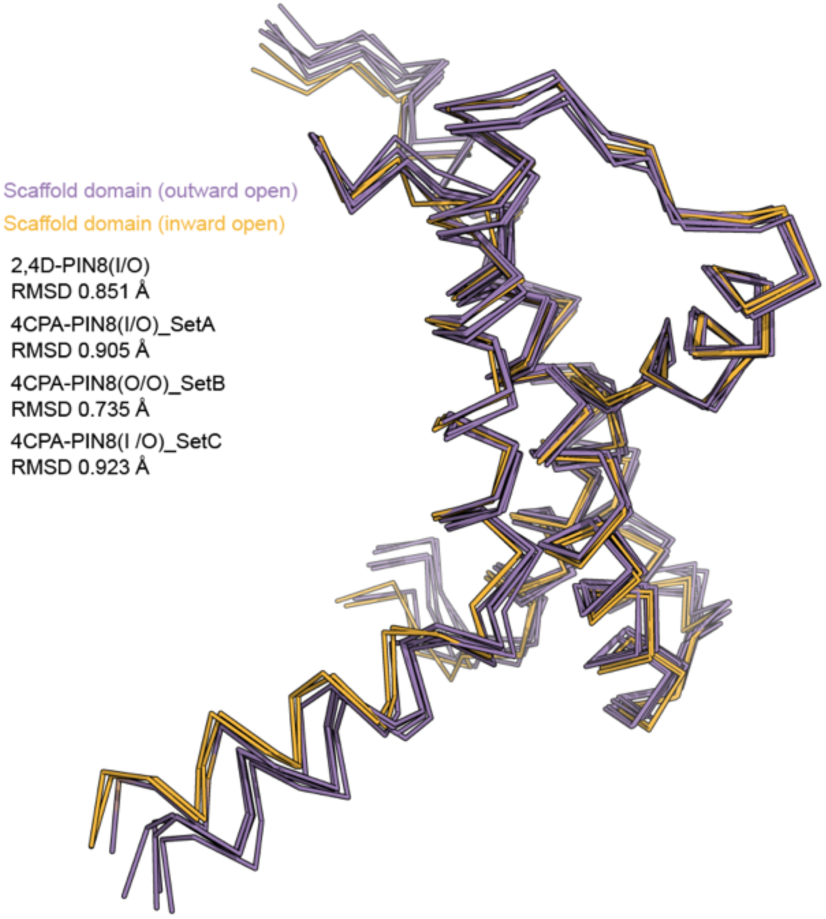
Scaffold domain during the transport. Superposition of scaffold domain in different transport states show that the scaffold domain does not change during transport.

**Extended Data Table 1:**
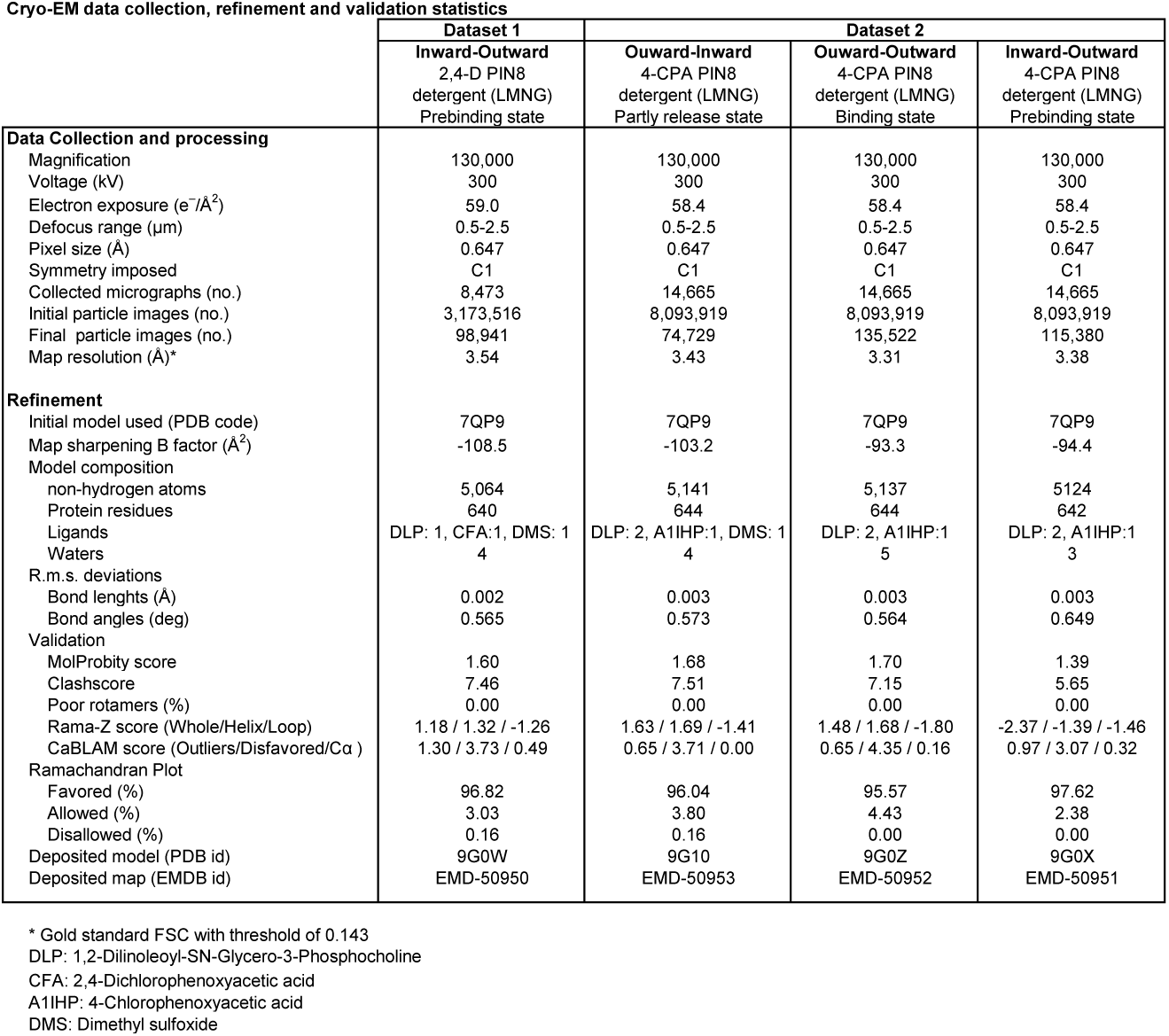
Statistics for cryo-EM data collection, model refinement and validation.

**Extended Data Table 2:** Results of the statistical analyses performed in Extended Data Fig. 1H and I. Groups were compared by a two-sided unpaired Student’s t-test.

| Substrate | NPA | Segment | P value |
| --- | --- | --- | --- |
| IAA | -NPA | WT<sup>Seg1</sup> vs *pin1*<sup>Seg1</sup> | 0.0005 |
|  |  | WT<sup>Seg2</sup> vs *pin1*<sup>Seg2</sup> | 0.001 |
|  |  | WT<sup>Seg3</sup> vs *pin1*<sup>Seg3</sup> | 0.036 |
|  | +NPA | WT<sup>Seg1</sup> vs *pin1*<sup>Seg1</sup> | 0.7145 |
|  |  | WT<sup>Seg2</sup> vs *pin1*<sup>Seg2</sup> | 0.8337 |
|  |  | WT<sup>Seg3</sup> vs *pin1*<sup>Seg3</sup> | 0.0909 |
| 2,4D | -NPA | WT<sup>Seg1</sup> vs *pin1*<sup>Seg1</sup> | 0.0042 |
|  |  | WT<sup>Seg2</sup> vs *pin1*<sup>Seg2</sup> | 0.0781 |
|  |  | WT<sup>Seg3</sup> vs *pin1*<sup>Seg3</sup> | > 0.9999 |
|  | +NPA | WT<sup>Seg1</sup> vs *pin1*<sup>Seg1</sup> | 0.1638 |
|  |  | WT<sup>Seg2</sup> vs *pin1*<sup>Seg2</sup> | 0.4574 |
|  |  | WT<sup>Seg3</sup> vs *pin1*<sup>Seg3</sup> | 0.4979 |

**Extended Data Table 3:** Results of the statistical analysis shown in Fig. 2A. One-way ANOVA followed by a Turkeýs multiple comparisons test (p < 0.05).

| Tukey's multiple comparisons test | P value |
| --- | --- |
| IAA vs. 2,4-D | >0.9999 |
| IAA vs. 4-CPA | <0.0001 |
| IAA vs. MCPA | 0.3977 |
| IAA vs. 2-CPA | 0.8412 |
| IAA vs. PA | <0.0001 |
| IAA vs. 2,4-DPP | >0.9999 |
| IAA vs. 2,4,5-T | 0.9995 |
| IAA vs. 2-MPA | 0.0013 |
| IAA vs. 4-MPA | <0.0001 |
| IAA vs. 2,4-DMPA | 0.0826 |
| 2,4-D vs. 4-CPA | 0.0002 |
| 2,4-D vs. MCPA | 0.6967 |
| 2,4-D vs. 2-CPA | 0.9808 |
| 2,4-D vs. PA | <0.0001 |
| 2,4-D vs. 2,4-DPP | 0.9962 |
| 2,4-D vs. 2,4,5-T | 0.9711 |
| 2,4-D vs. 2-MPA | 0.0041 |
| 2,4-D vs. 4-MPA | <0.0001 |
| 2,4-D vs. 2,4-DMPA | 0.2088 |
| 4-CPA vs. MCPA | 0.0156 |
| 4-CPA vs. 2-CPA | 0.0027 |
| 4-CPA vs. PA | 0.0056 |
| 4-CPA vs. 2,4-DPP | <0.0001 |
| 4-CPA vs. 2,4,5-T | <0.0001 |
| 4-CPA vs. 2-MPA | 0.9512 |
| 4-CPA vs. 4-MPA | <0.0001 |
| 4-CPA vs. 2,4-DMPA | 0.1058 |
| MCPA vs. 2-CPA | 0.9993 |
| MCPA vs. PA | <0.0001 |
| MCPA vs. 2,4-DPP | 0.2002 |
| MCPA vs. 2,4,5-T | 0.1158 |
| MCPA vs. 2-MPA | 0.2304 |
| MCPA vs. 4-MPA | <0.0001 |
| MCPA vs. 2,4-DMPA | 0.9969 |
| 2-CPA vs. PA | <0.0001 |
| 2-CPA vs. 2,4-DPP | 0.5929 |
| 2-CPA vs. 2,4,5-T | 0.4132 |
| 2-CPA vs. 2-MPA | 0.0538 |
| 2-CPA vs. 4-MPA | <0.0001 |
| 2-CPA vs. 2,4-DMPA | 0.8334 |
| PA vs. 2,4-DPP | <0.0001 |
| PA vs. 2,4,5-T | <0.0001 |
| PA vs. 2-MPA | 0.0003 |
| PA vs. 4-MPA | 0.1258 |
| PA vs. 2,4-DMPA | <0.0001 |
| 2,4-DPP vs. 2,4,5-T | >0.9999 |
| 2,4-DPP vs. 2-MPA | 0.0005 |
| 2,4-DPP vs. 4-MPA | <0.0001 |
| 2,4-DPP vs. 2,4-DMPA | 0.0333 |
| 2,4,5-T vs. 2-MPA | 0.0002 |
| 2,4,5-T vs. 4-MPA | <0.0001 |
| 2,4,5-T vs. 2,4-DMPA | 0.0173 |
| 2-MPA vs. 4-MPA | <0.0001 |
| 2-MPA vs. 2,4-DMPA | 0.7308 |
| 4-MPA vs. 2,4-DMPA | <0.0001 |

**Extended Data Table 4:** Statistics to support the Binding analyses shown in Extended Data Fig. 3. A Michaelis–Menten model is fit to describe kinetics.

| Substrate | Mutant | $r^2$ | $EC_{50}$ (μM) | $I_{max}$ (nA) |
| --- | --- | --- | --- | --- |
| IAA | WT | 0.9579 | $254 \pm 6$ | $18 \pm 3$ |
| | S146A | 0.8702 | $498 \pm 191$ | $8 \pm 2$ |
| | Y150F | 0.8601 | $452 \pm 32$ | $6 \pm 2$ |
| | Y150A | 0.9395 | $1206 \pm 298$ | $4 \pm 0.3$ |
| | I51Y | 0.8627 | $547 \pm 171$ | $2 \pm 0.2$ |
| | S266A | 0.8679 | $578 \pm 183$ | $4 \pm 0.6$ |
| | N223A | 0.8533 | $444 \pm 205$ | $2 \pm 0.8$ |
| | S55A | 0.8628 | $213 \pm 58$ | $6 \pm 1$ |
| | S55A/S146A | 0.7788 | $341 \pm 128$ | $7 \pm 2$ |
| 4-CPA | WT | 0.9556 | $800 \pm 176$ | $12 \pm 2$ |
| | S146A | 0.9073 | $369 \pm 27$ | $8 \pm 2$ |
| | Y150F | 0.9167 | $710 \pm 228$ | $5 \pm 0.9$ |
| | Y150A | 0.9942 | $2616 \pm 263$ | $5 \pm 0.4$ |
| | I51Y | 0.9304 | $1044 \pm 252$ | $3 \pm 0.8$ |
| | S266A | 0.8493 | $2103 \pm 290$ | $7 \pm 2$ |
|  | N223A | - | - | - |
| | S55A | 0.9781 | $328 \pm 15$ | $11 \pm 1$ |
| | S55A/S146A | 0.9699 | $81 \pm 3$ | $11 \pm 0.9$ |
| 2,4D | WT | 0.9802 | $179 \pm 10$ | $17 \pm 2$ |
| | S146A | 0.9476 | $164 \pm 67$ | $8 \pm 0.9$ |
| | Y150F | 0.9703 | $816 \pm 275$ | $8 \pm 2$ |
| | Y150A | 0.9853 | $684 \pm 256$ | $5 \pm 1$ |
| | I51Y | 0.9281 | $641 \pm 77$ | $3 \pm 0.8$ |
| | S266A | 0.9800 | $774 \pm 70$ | $12 \pm 0.7$ |
|  | N223A | - | - | - |
| | S55A | 0.9282 | $95 \pm 36$ | $6 \pm 0.6$ |
| | S55A/S146A | 0.8915 | $38 \pm 6$ | $9 \pm 2$ |

